# Morphological profiling by high-throughput single-cell biophysical fractometry

**DOI:** 10.1101/2022.05.24.493226

**Authors:** Ziqi Zhang, Kelvin C. M. Lee, Dickson M. D. Siu, Queenie T. K. Lai, Edmund Y. Lam, Kevin K. Tsia

**Affiliations:** Department of Electrical and Electronic Engineering, The University of Hong Kong, Pokfulam Road, Hong Kong; Advanced Biomedical Instrumentation Centre, Hong Kong Science Park, Shatin, New Territories, Hong Kong

## Abstract

Complex and irregular cell architecture is known to statistically exhibit fractal geometry, i.e., a pattern resembles a smaller part of itself. Although fractal variations in cells are proven to be closely associated with the disease-related phenotypes that are otherwise obscured in the standard cell-based assays, fractal analysis with single-cell precision remains largely unexplored. To close this gap, here we develop an image-based approach that quantifies a multitude of single-cell biophysical fractal-related properties at subcellular resolution. Taking together with its high-throughput single-cell imaging performance (~10,000 cells/sec), this technique, termed single-cell biophysical fractometry, offers sufficient statistical power for delineating the cellular heterogeneity, in the context of classification of lung-cancer cell subtypes and tracking of cell-cycle progression. Further correlative fractal analysis shows that single-cell biophysical fractometry can enrich the standard morphological profiling depth and spearhead systematic fractal analysis of how cell morphology encodes cellular health and pathological conditions.

## Introduction

Cell morphology is constituted by the complex biomolecular machinery at the genomic, transcriptomics, and proteomic levels. Hence, it is a valuable readout, which can be captured by microscopy, for assaying the functional state of individual cells. Dramatic advancements in high-throughput imaging and computer vision in the past decade have sparked the major drive in the use of microscopy to extract quantifiable information from cell morphology (i.e., morphological profiling) ^1,2^. Creating the catalogues of the cell morphological features, this cell profiling strategy enables mining the underlying feature signatures or patterns that can infer cell age ^3^, metastatic potential ^4^, screening chemical ^5^ and genetic perturbation ^6^.

In morphological profiling, a wealth of quantitative metrics (features) can be extracted from the individual cell images, including cell size, shape, texture etc., representing a fingerprint of each cell. Downstream analysis is then applied to investigate the similarities or correlations between profiles in order to identify the phenotypes specific to the cell types and states. Traditionally, the morphological features are defined based on Euclidean geometry, which can be easily coupled with general variations in geometry (e.g., size and shape), however, irregular spatial information hidden in the complex cellular structure (e.g., statistical properties of shape and texture) could often be missed. This is particularly relevant to the cellular malignancy, in which the intracellular mass growth shows a significant degree of randomness and disorder ^7^. Specifically, conventional Euclidean geometry fails to holistically quantify the textural or shape irregularity at different length scales. This explains the need for an extended set of local and global features to separately examine the heterogeneity at different spatial scales ^8^. Yet, they do not capture an important property shared in a wide variety of biological cells, i.e., “fractality”. It refers to the fact that the texture/shape of an object does not significantly differ from the same property measured on the larger scale. To this end, fractal dimension (FD) has been adopted as an effective metric that quantifies and classifies the irregular biological structures and the self-similarity characteristics that are not well represented by the Euclidean geometry ^9^. FD is typically a non-integer value, in contrast to the dimensions defined in Euclidean geometry, i.e., 1 for a line (1D), 2 for a plane (2D) and 3 for a cube (3D).

Indeed, fractal analysis has been demonstrated an effective tool in clinical diagnosis, such as examination of aberrant histopathological features in tissues ^10^, and assessment of abnormal organ morphology (e.g. tumor vasculatures) in radiology ^11^. Further down to the cellular or even subcellular level, fractal behavior can be observed in the chromatin topology in the nucleus ^12–14^, cell membrane contour and adhesion topology ^15^, mitochondrial ^16^ and cytoskeleton ^17^ morphology. For instance, the mitochondria organization undergoing fission and fusion regulated by cellular metabolism follows the statistics of self-similarity ^18^; the protein interaction and structural interminglement of chromatin are both highly consistent with a fractal framework ^12^; the architecture of cytoskeleton and plasma membrane also shows a self-similar topology where molecules are confined hierarchically over time and length scales ^19^. Hence, the knowledge of fractal characteristics of cells could offer new physical insights of cell types and states. Specifically, changes in cellular and subcellular FDs are now known to be closely related to the epigenetic states ^20^, the gene expression levels ^16^. Hence, they can be indicative of the functional states of cells (e.g., metabolic states ^21^, cell differentiation states ^22^, and cell malignancy ^23^).

However, these promises, together with the cellular fractal characteristics that can readily be analyzed by standard microscopy, have not yet made fractal analysis widely applicable in cytometry and morphological profiling of cells. The key challenge stems from the fact that the cellular/subcellular morphology exhibits fractal properties in a statistical sense, instead of the archetypal geometrical sense. Yet, current imaging techniques lack the scale and throughput to guarantee that fractal analysis could show the sufficient statistical power for delineating the cellular heterogeneity and complexity on the single-cell level (e.g., limited to ~10’s −100’s single cells ^15,23,24^).

To address this challenge, here we employ an ultrahigh-throughput quantitative phase imaging (QPI) flow cytometer called multiplexed asymmetric-detection time-stretch optical microscopy (multi-ATOM) ^25,26^ to analyse single-cell biophysical fractal characteristics (termed *single-cell biophysical fractometry*) at the breadth and depth not achievable by the existing methods. This strategy of single-cell fractometry is achieved by two key attributes: (1) Establishing label-free morphological profiling that includes not only the common shape and texture features based on Euclidean geometry, but also a collection of biophysical fractal parameters (not only FD) of each cell. This is enabled by the core strength of QPI inherited by multi-ATOM, in which the *complex-field* image information of individual cells (i.e., both amplitude and quantitative phase images) can be obtained at subcellular resolution. Such complex-field information can then be harnessed to compute the corresponding far-field light scattering pattern (by means of Fourier Transform light scattering (FTLS) ^27^), which provides a catalogue of single-cell fractal and the associated ALS features. Defining these fractal features as an intrinsic morphological profile aligns precisely with the growing interest in new strategies for in-depth biophysical phenotyping of cells, that has already generated new mechanistic knowledge of cell heterogeneity and showed initial promises in identifying cost-effective biomarkers of disease, thanks to its label-free nature ^28^. (2) Enabling large-scale single-cell fractometry by the ultrafast QPI operation in multi-ATOM, at the speed at least 100 times faster than the existing QPI modalities that rely on camera technology for image recording ^29^. Combined with the high-throughput microfluidics platform ^8,25,26,30^, this approach enables single-cell fractometry at a throughput of at least 10,000 cells/sec without sacrificing the subcellular imaging resolution. This attribute critically provides in-depth statistical fractal analysis, which has largely been underexploited in the previous work on fractal analysis, especially at the single-cell screening resolution. Indeed, comprehensive morphological profiling often relies on analysing the deeper statistics of cell phenotypes in order to detect cellular heterogeneity and subpopulation with an improved sensitivity and robustness ^31^.

In this work, we show that high-throughput single-cell biophysical fractometry allows us to distinguish the histologically differentiated subtypes of lung cancer cell by fractal-related features. We also demonstrate that these fractal-related features play an important role in identifying different stages of cell cycle progression (G1, S and G2). To gain a better interpretation of the significance of the extracted fractal-related features, we further investigate the underlying connections, if any, between Euclidean-defined morphological features and the fractal features.

## Results

### Key workflow and basic performance of single-cell biophysical fractometry

We applied multi-ATOM, an ultrafast QPI modality, to perform high-throughput single-cell imaging in microfluidic flow (See **Methods**) (**Fig. 1a**). Detailed working principle and experimental configuration were reported previously ^8,25,26^. The general principle is to first record the complex optical field at the image plane of the flowing cell by multi-ATOM, i.e., *E*(*x*, *y*) = *A*(*x*, *y*)e^*jϕ(x, y)*^ where *A*(*x, y*) is amplitude (i.e. bright-field) image and *ϕ*(*x, y*) is the quantitative phase image (**Fig. 1b**). Subsequently, the complex field at the image plane is then numerically propagated to the far field using the Fourier transform operation – yielding the (far-field) scattered light-field pattern 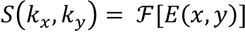^27^, from which the fractal properties of the cell can be measured (**Fig. 1c**). It is due to the fact that the cellular and subcellular fractal structures give rise to the heterogeneity of the refractive index within the cells, and thus directly impact the scattered light pattern ^32^. Here, we further convert the scattered light pattern into an angular light scattering (ALS) profile *S*(*q*) in which scattered light intensity is averaged over rings of constant wave vector *q* = 4*π*/*λ sin* (*θ*/2), where *θ* is the *polar* scattering angle ^27^ (**Fig. 1d**). This approach has been adopted in characterizing different metabolic states of red blood cells (RBCs) ^33^, assessing different intracellular organelles ^34^, and classification of bacterial species ^35^, and analyzing the fractal characterization of the fibrin network ^36^, all in a label-free manner.

**Figure 1.**
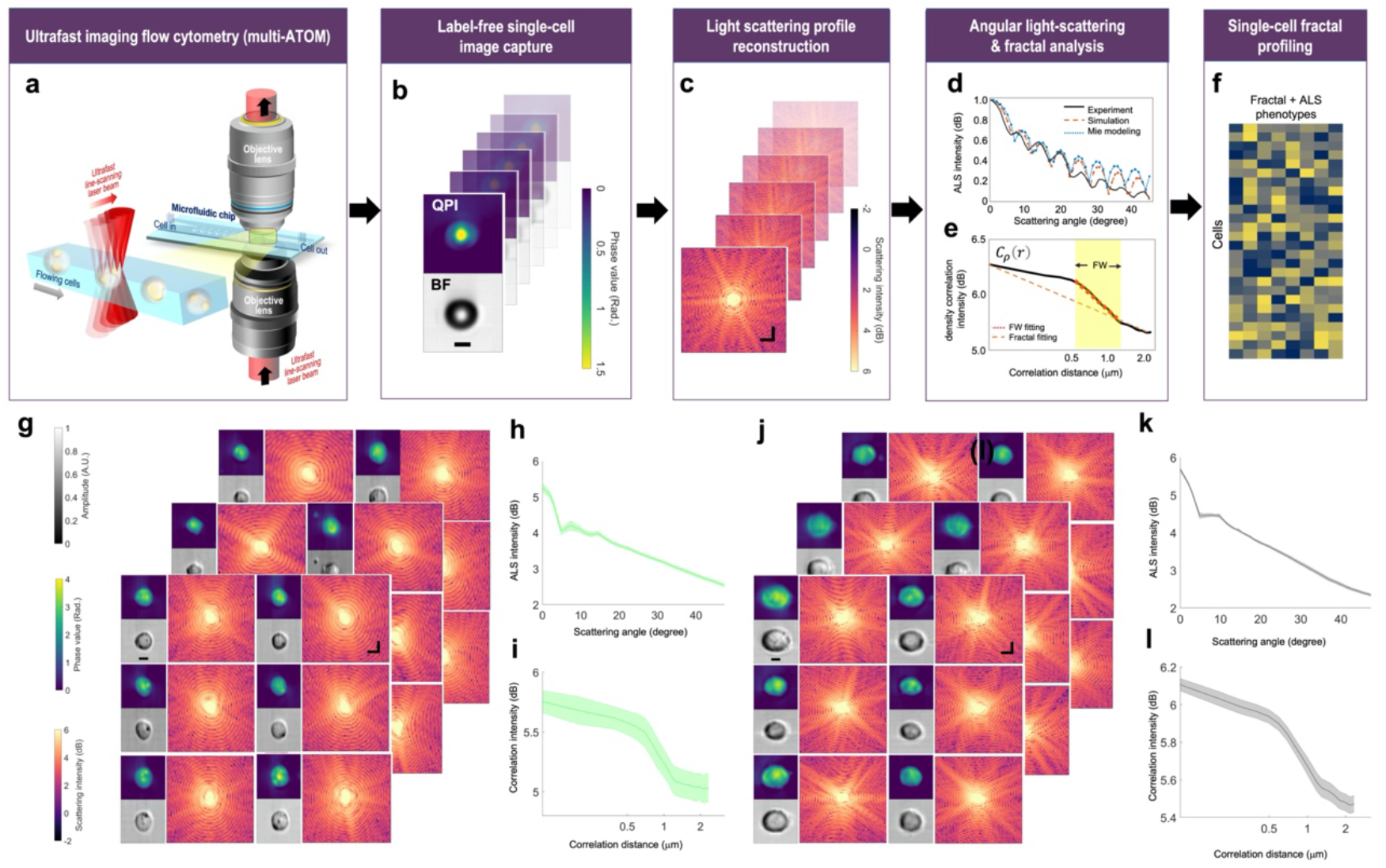
General workflow and performance of high-throughput single-cell fractal profiling. **(a)** Ultrafast imaging flow cytometry by multi-ATOM. **(b)** Label-free single-cell image capture (Top: QPI; bottom: Bright-field image (BF)) (Example: polymer microspheres). Scale bar = 5 μm. (**c)** Complex-field light scattering profile reconstruction via FTLS (Example: polymer microspheres). Scale bar = 2 rad/μm. **(d)** Retrieval of the ALS profile. (Example: Experimental microsphere data acquired by multi-ATOM (red) and theoretical simulation result (blue) and the analytical Mie light scattering theory of ideal sphere with the same size as the spheres (black). **(e)** Schematics of fractal fitting of *C_ρ_*(*r*). FD is determined from the fractal fitting of the overall curve, whereas the fitting within FW gives FD with FW. **(f)** ALS and fractal profiles extracted from **(d) and (e)**. **(g)** Label-free single-cell image capture of leukemia cell line (ACC220). Spatial scale bar = 5 μm. Frequency scale bar = 2 rad/μm. (**h**) ALS plots summarizing ALS profiles of 2,500 ACC220 cells. Shaded area indicates the statistical variance. **(i)** Plot summarizing *C_ρ_*(*r*) of 2,500 ACC220 cells. Shaded area indicates the statistical variance. **(j)** Label-free single-cell image capture exampled by leukemia cell line (THP-1). Spatial scale bar = 5 μm. Frequency scale bar = 2 rad/μm. (**k)** ALS plot summarizing ALS profiles of 2,500 THP-1 cells. Shaded area indicates the statistical variance. **(l)** Plot summarizing *C_ρ_*(*r*) of 2,500 THP-1 cells. Shaded area indicates the statistical variance.

Based on the light scattering theory ^37^, we can further relate the ALS with the dry-mass density variation *ρ*(*r*) (quantified through spatial correlation of the density fluctuation *C_ρ_*(*r*)) (See **Methods**) ^38^ in such a way that the Fourier transform of an ALS intensity profile will obey an inverse power law relationship, i.e., 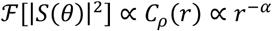, where *α* is the exponent, and *C_ρ_*(*r*) is the spatial correlation of *ρ*(*r*). In practice, by fitting the slope *α* of the log-scaled plot of 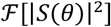, we could calculate the FD = 3 – *α* (See detailed derivation in **Methods**) (**Fig. 1e**). As cells exhibit fractal properties (e.g., self-similarity) only within a limited range of length scales, referred to as the fractal window (FW), ALS offers an effective tool to identify the FW where the inverse power law behavior is present ^21,39^. In addition, as mentioned earlier, the fractal behaviors of cells are manifested in a quantitative statistical sense. We exploited a catalogue of parameters that quantify the statistics of the ALS profiles, and the statistics related to FW fitting, e.g., the mean square error (MSE), the FW width and the estimated FD (See the complete list of parameters in **Supplementary Table S1**) (**Fig. 1f**). We note that these FD-related parameters can reflect how well the fractality is preserved in different scales and allow us to quantify the degree of self-similarity. The basic performance of imaging and the ALS analysis were tested with polystyrene microbeads (**Fig. 1b-d**), and two different types of leukemic cells, ACC220 (**Fig. 1g-i**) and THP-1 (**Fig. 1j-l**).

### Single-cell fractal profiles distinguish different lung cancer cell subtypes

Cell morphology assessment is commonly practiced in cancer diagnosis and classification. Taking lung cancer, the leading cause of cancer-related mortality worldwide ^40^, as an example, histological characterizations of small biopsies or cytology specimens play an integral role in the pipeline for classifying different lung cancer types, according to the criteria long-established by World Health Organization (WHO) ^41,42^. However, these assessments are often confounded by subjective and biased visual inspection and are mostly limited to obvious morphological abnormalities across the histochemically stained tissues, e.g., cell shapes and intercellular textural complexity. As abnormal subcellular morphology (e.g. nucleus/nucleoli and cytoplasm) is also found to be indicative of malignancy and different cancer subtypes ^43,44^, we sought to investigate if the single-cell fractal properties of different lung cancer cell subtypes extracted from the label-free ALS profiles can provide unbiased classification of the three key histologically different lung cancer subtypes: small cell lung carcinoma (SCLC) and two subtypes of non–small cell lung carcinoma (NSCLC), which are squamous cell carcinoma (SCC) and adenocarcinoma (ADC) ^41^.

Based on the visual examination of the randomly selected reconstructed single-cell phase gradient and quantitative phase images captured by multi-ATOM (**Fig. 2a**), it is generally impractical to distinguish the 3 subtypes by standard bulk features such as cell size and optical density (or opacity). Notably, although the general size distribution of the population differs between SCLC and NSCLC, there is a significant size variation on the single-cell level even within the same subtype because of the heterogeneity among individual cells. Therefore, cell size alone is not an effective feature for single-cell identification between SCLC and NSCLC in such as broad populations. Going beyond these bulk features, we evaluated the *C_ρ_*(*r*) from the ALS profile (*S*(*θ*)) of individual cells (**Fig. 2a**). We observed that *C_ρ_*(*r*) appears to be statistically different among 3 subtypes (including a total of 7 different lung cancer cell lines (**Supplementary Fig. S1**)). Furthermore, we calculated the FD of individual cells through inverse power law fitting and observed that the FD distributions can relatively be categorized into the low/middle/high level for SCLC, SCC and ADC, respectively (**Fig. 2b**), with a significant effect size (|*d*| ~0.57-0.89 between subtypes using Cliff’s delta statistics). The statistics of the overall fitting error (FD MSE2) also shows significant difference among the subtypes (|*d*|> 0.57 – 0.88). More importantly, FD MSE2 indicates that there is a larger variance (or dispersion) of *C_ρ_*(*r*) for SCLC, while the cells of SCC and ADC both tend to keep a better linearity, implying the self-similarity of cellular structure is more consistently preserved in a wider length scale (**Fig. 2c**). We stress that effect size, which is independent of sample size, is adopted here to show the significance of sample difference. This is due to the fact that the common *p*-value will show misleading high statistical significance when using a large sample size ^45^, which is challenging to achieve in other fractal cellular measurements but realized by our multi-ATOM system (> 10,000 cells). Besides effective size analysis, we also computed the Spearman correlation coefficient of fractal features and lung cancer subtype (1 for SCLC, 2 for SCC and 3 for ADC) to prove their close bonding (FD: 0.7174; FD MSE2: −0.6791).

**Figure 2.**
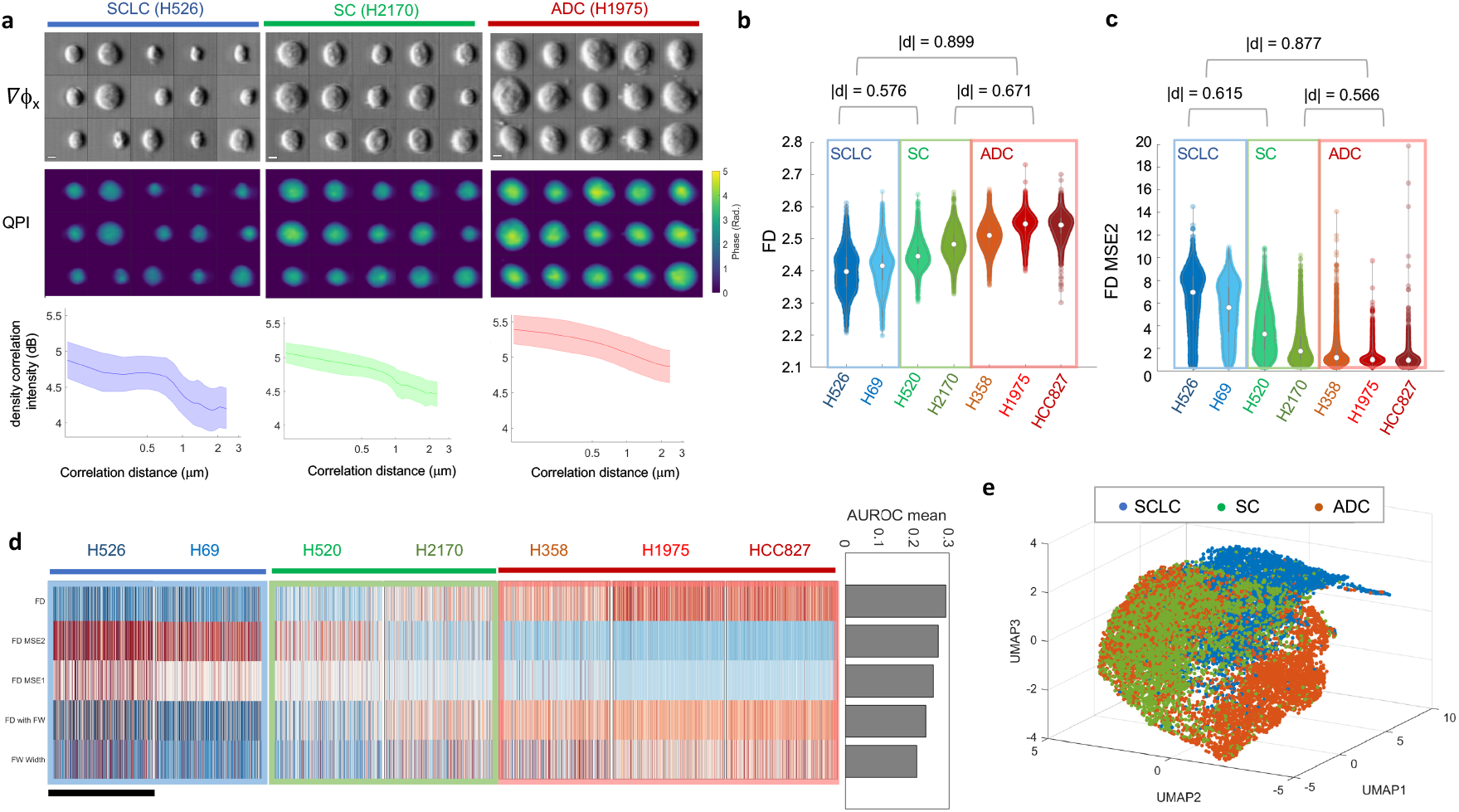
Fractometry of multiple lung cancer cell subtypes. **(a)** Randomly selected phase gradient images (Δ_ϕx_, top), QPI (middle) and the correlation function plots (bottom) of 3 lung cancer cell lines, respectively. Scalar bar is 5 μm. Shaded area indicates the statistical variance. **(b)** Statistical distributions of FD. **(c)** Statistical distributions of FD MSE2. **(d)** A fractal phenotypic profile of 7 lung cancer cell lines. Each row represents a fractal feature, and each column represents a single cell. The scale bar stands for 1,000 cells, which were randomly subsampled from each cell line. The AUROC ranking results are shown in the right panel. **(e)** 3D UMAP visualization of the 17 fractal and ALS-related phenotypes extracted from single-cell images of the lung cancer cell lines. Clusters are colored according to the three lung cancer subtypes.

Leveraging the statistical power offered by multi-ATOM, we further extracted the statistics of other fractal-related features from *C_ρ_*(*r*) (normalized based on the *z*-score, subsampled from randomly selected 1,000 cells per cell line) to form a fractal profile for each cell (**Fig. 2d**). Based on the heatmap of the five most significant fractal features (ranked by area-under-curve of the receiver operating characteristics (AUROC) of one-versus-all classification (right panel of **Fig. 2d**), we observe that each of the three lung cancer subtypes exhibits its distinct characteristic pattern in the fractal profile. For instance, the three ADC cell lines (H358, H1975, HCC827) share the similar profile that shows high FD, low FD MSE2 and high FD width. When we further included other ALS features for higher-dimensional analysis (a total of 17 dimensions, see **Supplementary Table S1**), visualized by the uniform manifold approximation and projection (UMAP) algorithm (**Fig. 2e**), we observed the three distinct main clusters corresponding to the key lung-cancer subtypes. Although overlapping is observable to some degree in the clusters of ADC and SCC, most cells of SCLC are highly dispersed from NSCLC. Hence, this study suggests that morphological profiling based on these label-free single-cell fractal features, which are closely linked to the intracellular mass-density distribution characteristics, could provide the discriminative power to distinguish the key histologically different lung cancer subtypes.

### Single-cell fractal profile recapitulates cell-cycle progression

Going beyond cell-type classification, we sought to investigate if and how the label-free single-cell fractal characteristics are impacted by different cell states in the cell-cycle progression. In this study, the single-cell QPI/BF image recording of over 15,000 fixed breast cancer cells (MDA-MB231) is synchronized with 2-color fluorescence detection (**Fig. 3a**, and see **Methods**). This multimodality allows us to correlate the label-free fractal properties with the DNA content quantified by the fluorescence labels, at the single-cell precision, as the ground truth of cell-cycle progression from G1, S to G2 phase (by the propidium iodide (PI) label which quantifies DNA content whereas EdU (5-ethynyl-2’-deoxyuridine) indicates the newly synthesized DNA in S-phase cells) (**Fig. 3b**). We note that the variation in the DNA content, and thus the changes in the biophysical properties (including the fractal characteristics) reflects the continuous progression of cells, instead of discrete states of G1, S, and G2 ^46^. Hence, the “ground truth” given by the 2-color cell-cycle fluorescence markers/labels should cautiously be treated as the reference, which allows us to interpret the biophysical properties (especially FD) based on the established biochemical signatures (e.g., DNA synthesis and replication) and the related biological events (e.g., cell growth and protein synthesis).

**Figure 3.**
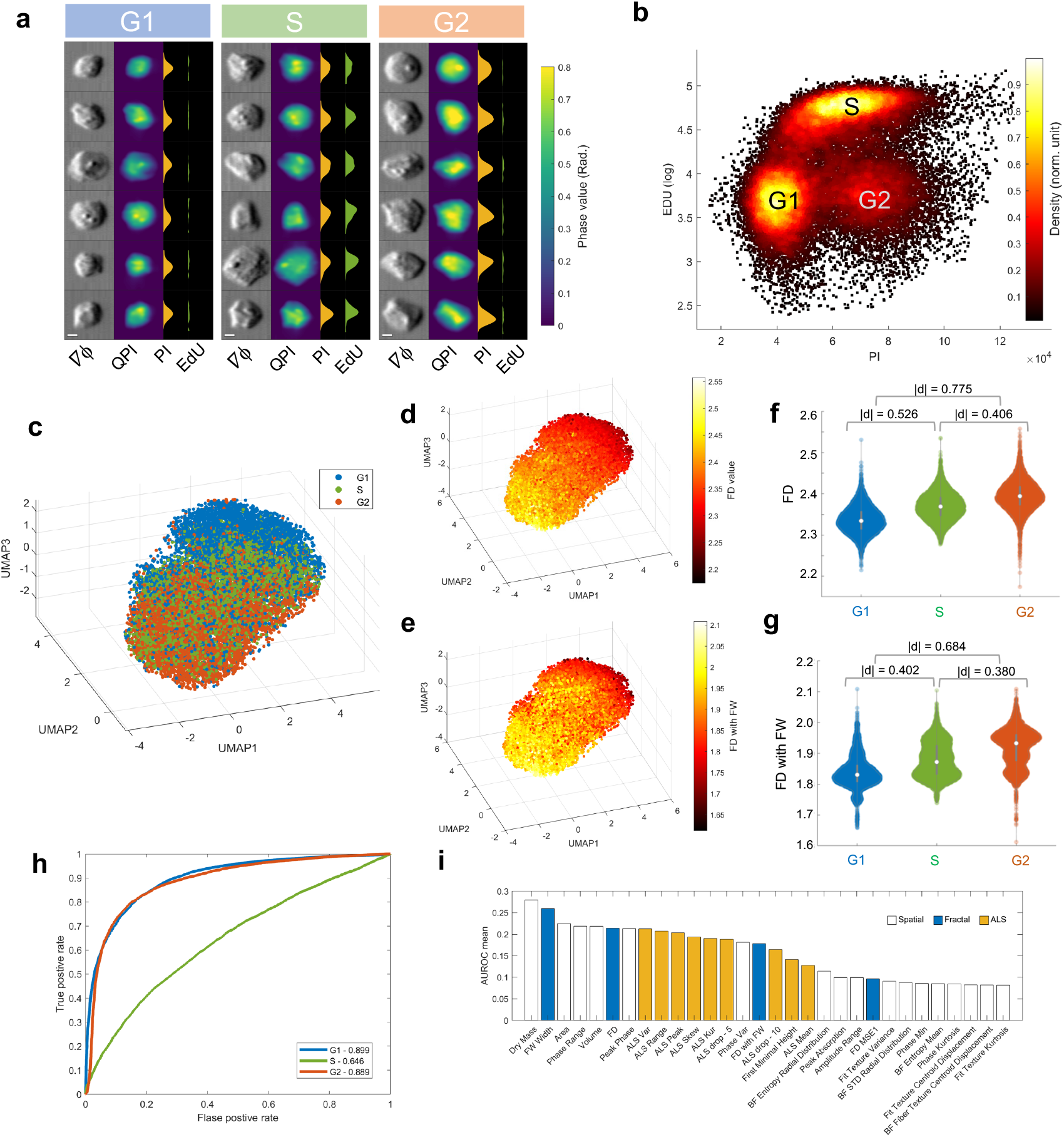
Fractal analysis of cell cycle progression (MB231 cells). **(a)** Randomly selected single-cell images (Phase gradient (Δ*ϕ*) and QPI) and the synchronized fluorescence detection of the same cells in different cell cycle phases (G1, S, G2): (From left to right), Δ*ϕ* images, QPI images, fluorescence profile of PI, and fluorescence profile of EdU. Scale bar = 5 μm. **(b)** Synchronized 2-color (EdU versus PI) fluorescence detection in multi-ATOM. DNA content is quantified by PI intensity, whereas the S-phase cells are recognized by EdU (in log scale). A standard flow cytometry result for cell cycle determination is shown in **Supplementary Fig. S3** for reference. **(c)** 3D UMAP visualization of the full set of phenotypes (both Fourier and spatial) showing the trend of cell cycle progression. **(d,e)** The same UMAP plot in **(c)** color-coded with the FD value, the FW value, respectively. **(f,g)** Violin plots of FD, and FD with FW, respectively, across the G1, S, and G2 phase. **(h)** ROC curves of the linear regression classifier constructed from all the fractal and ALS features. AUROC values are labeled in the legend. **(i)** Feature ranking by AUROC mean (Top 30 features).

To further harness the strength of information-rich morphological profiling, we defined an extensive set of multi-faceted label-free morphological readouts, encompassing the spatial features (directly computed from QPI and BF following a hierarchical strategy ^8^), the light-scattering characteristics (extracted from ALS profile), as well as the fractal profile. We observed that this label-free profile revealed the overall trajectory of the cell-cycle progression from G1, S to G2 phase in the UMAP visualization (**Fig. 3c**). Importantly, the expression variations of the key fractal features, such as FD and FD with FW, also consistently follow the progression (**Fig. 3d-e**), and show significant differences across the three phases (|*d*| > 0.40 for FD; |*d*| > 0.38 for FD with FW) (**Fig. 3f-g**). FD exhibits a progressive increase along the G1-S-G2 order (**Fig. 3f)**, which is consistent with the trend shown by cell size and cell mass (**Supplementary Fig. S2**), as the size enlargement and mass accumulation are common biophysical traits during cell cycle progression. This suggests a growing complexity and irregularity of intracellular mass distribution, which could be attributable to DNA replication and the subsequent protein synthesis process (e.g. microtubule production) as the cell evolves from the G1, S to G2 phase.

We further assessed the ability of using fractal features to identify the three cell-cycle phases in a one-versus-all mode through the receiver operating characteristic (ROC) analysis. While using FD alone is found to be not sensitive to detect the S-phase, this single fractal feature is on the other hand effective in identifying the G1 and G2 phases, with the AUROC of 0.822 and 0.790 for G1 and G2 phase, respectively. As S phase is a transit state between G1 and G2, it is acceptable that more features are needed for an accurate identification. Therefore, we performed the same ROC analysis with a linear regression classifier integrating all the features extracted from Fourier domain (17 dimensions), and found an improvement of the AUROC for all the three phases (G1:0.899, S:0.646, G2:0.889), especially for S phase (**Fig. 3h**).

By performing the same ROC analysis including all the features extracted from QPI/BF morphology the ALS features and the fractal profile (**Supplementary Fig. S4**), we also quantified the significance of these features in performing the (one-versus-all) classification of the cell-cycle phases (**Fig. 3i**). We observed that, apart from the cell size and cell mass, which are known to be tightly linked to cell-cycle progression, multiple fractal and ALS features (e.g., FW width and FD) extracted from the FTLS analysis are among the top 30 features with the highest averaged AUROC, which implicates the informativeness of these Fourier-domain features.

Notably, the high-rank of FW width could suggest that an irregular cell mass growth and distribution occur across a longer length-scale (i.e., the scale-invariant property of fractals). We further visualized the expression pattern of a total of 101 morphological features (see the circular heatmap in **Fig. 4a**) and observed that more than half of the spatial features extracted by Euclidean geometry (from the bulk, global to local spatial features) do not show as clear changes as many ALS and fractal features across the three cell-cycle phases. The above analyses suggest that the fractal features could offer the label-free specificity and sensitivity to track the cell-cycle progression.

**Figure 4.**
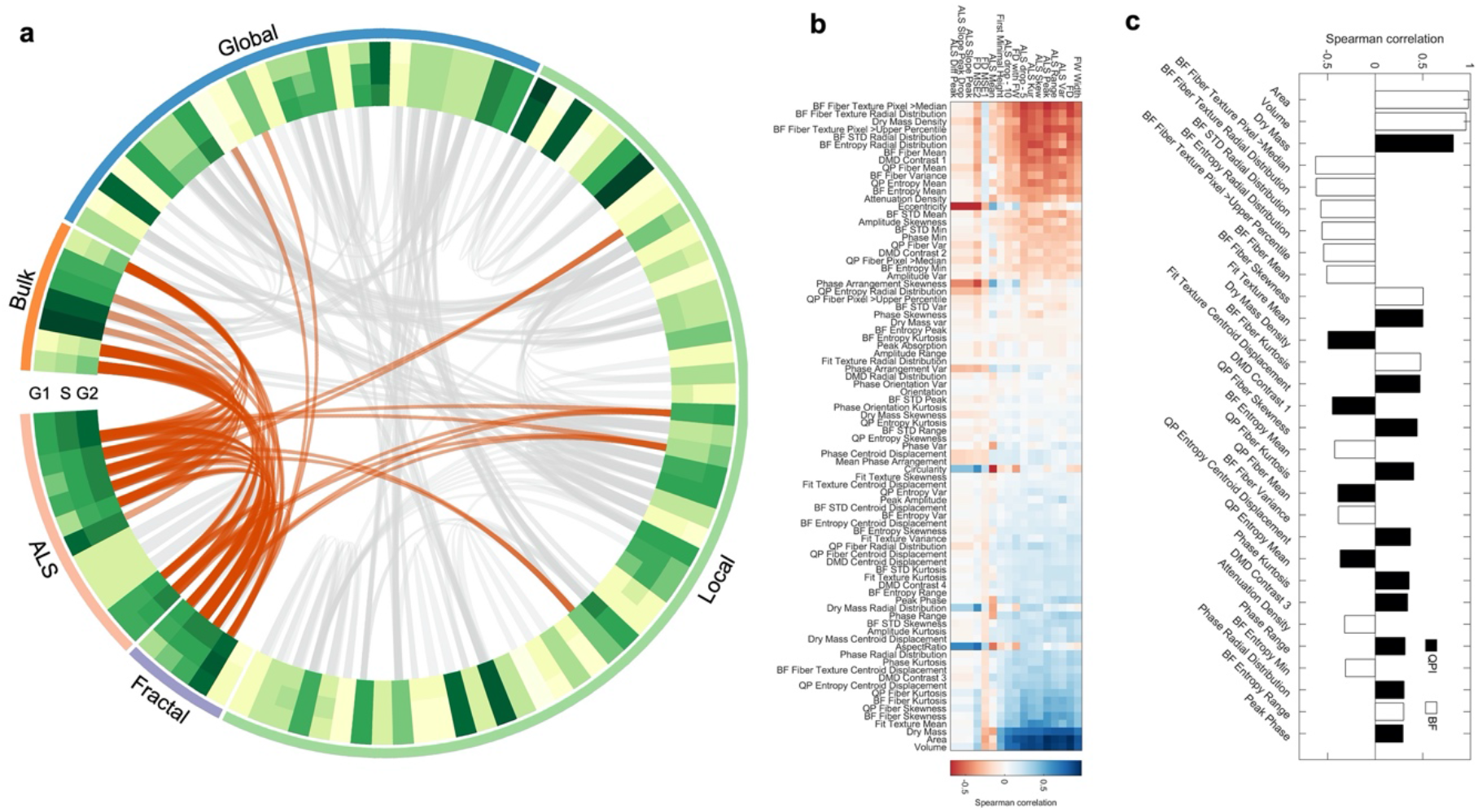
Analysis of the single-cell phenotypic correlation among the fractal, ALS and spatial features. **(a)** A Circular plot summarizing the mean heatmap and the correlations among all features in cell-cycle progression. The feature type is labelled by the outmost colored ring (bulk - orange, global - blue, local - green, fractal – purple and ALS - pink), and the full labels can be referred to **Supplementary Fig. S6**. The mean feature values of in different cell cycle phases are color-coded in the three ring-shaped heatmap. In the inner circle, all the feature pair with an absolute value of Spearman correlation coefficient over 0.6 are linked together by gray lines, while the Fourier-morphology connections are colored with orange specifically. Thickness of the lines is also encoded by the absolute value of correlation coefficient. **(b)** Spearman correlation heatmap between Fourier-domain and spatial-domain features. Fourier features are ranked by AUROC test. Spatial domain features are ranked by the average of correlation coefficient with all Fourier features. **(c)** Correlation coefficient bar chart of FD ranked by absolute value. Only 30 features with the largest coefficient magnitude with FD are listed.

We further investigated if and how the common spatial features extracted by Euclidean geometry can be correlated with the ALS and fractal features during the cell cycle progression (a total of 101 dimensions) – gaining additional insight of these classical morphological features in the context of the fractal behavior (**Fig. 4a-c**). We observed that the Fourier features, i.e., ALS and fractal features, are in general not strongly correlated with the (Euclidean) spatial features, except a handful of 12 spatial features (i.e., ~12% of all the features) with the absolute value of Spearman correlation coefficients > 0.6, see the orange lines in **Fig. 4a and the heatmap in Fig. 4b**). It indicates the low redundancy between these two classes of features in describing the morphological characteristics of cells. We noted that the fractal and some ALS features are particularly correlated with the cell (dry) mass and size – all of which are the important features sensitive to the cell cycle progression, consistent with the earlier feature ranking analysis (**Fig. 3i**). We further identified that the ALS features are more favorably linked to the local textures (derived from both the BF and QPI images) (**Fig. 4b**). For instance, by ranking the spatial features according to their correlations with FD (**Fig. 4(c)**), we further identified a tight connection between the changes in FD and dry mass, the local textures that are related to the statistics of the fiber textures and structural entropy. Based on this analysis, we thus could enrich the label-free morphological profile that characterizes the biophysics of cell cycle progression. Not only the well-known features of cell and mass varies during the cell cycle, but also the local textures linked to subcellular mass density (from QPI) and optical density (from BF) distributions, many of which follow the fractal behavior. A full view of the correlation analysis can be referred to **Supplementary Fig. S5**.

## Discussion

Fractal characteristics of cell morphology has been well acknowledged for four decades ^47^, and has also been proven indicative of complex cellular functions, especially disease progression. However, practices of defining cellular fractal phenotypes have not been widely adopted in cytometry and cell-based assay. This gap stems from that the fractal feature, e.g., FD, is a statistical measure reflecting the morphological structural complexity (especially the self-similar properties) of cellular/sub-cellular components. Yet, current imaging cell-based assays lack the throughput to demonstrate the fractal analysis with sufficient statistical power for delineating cellular heterogeneity, especially at the single-cell precision. In this study, we demonstrated a high-throughput morphological profiling strategy (~10,000 cells/sec) that empowers biophysical single-cell fractometry based on a catalogue of label-free single-cell fractal-related features extracted from an ultrafast QPI flow cytometry platform. Specifically, the information content of the single-cell fractal profile is enriched by (1) harnessing the fractal signature of a cell is intimately linked to its light scattering characteristics, which can readily be read out by our QPI platform through *single-cell* FTLS analysis; and (2) quantifying the statistics of these single-cell fractal features, thanks to the throughput offered by the QPI platform in this work.

We stress that QPI, in contrast to other optical imaging modalities, has a unique capability of quantifying dry mass density distribution of a cell at high sensitivity, that is derived from the optical path length profile given by QPI (**Methods**) ^48^. For instance, in the current multi-ATOM platform, the detection sensitivity of the optical path length was reported to be as small as 4 – 8 nm ^25^, which corresponds to the dry mass surface density resolution of ~ 0.02-0.04 pg/mm^2^, taking the refractive increment of 0.19 ml/g for biological cells ^49^. Therefore, biophysical single-cell fractometry described in this work measures not only the fractal properties of the cellular morphology with subcellular resolution, but also the subtle variation in the *(dry) mass* fractal characteristics.

We demonstrated that the label-free single-cell fractal profiles exhibited the discriminative power for unbiased classification of the histologically different lung cancer subtypes, and are highly correlated with the cell-cycle progression. More importantly, the fractal profile could also be integrated with the conventional morphological profile based on the spatial features extracted using the Euclidean geometry. This enables extensive feature extraction which has a two-fold benefit: first, it further augments the profiling dimensionality (and thus potentially better encode relevant biological information). This allows us to identify feature correlation, based on which further feature selection (e.g., removing noisy or redundant features ^50^) can be done for extracting/refining relevant information for downstream analysis. Second, it permits us to mine the potential correlative patterns between fractal features and other spatial Euclidean features. Hence, we could gain better interpretability, and thus biophysical insights of the morphological features, in the context of fractal behaviour (e.g., the correlation between FD and the local fiber textures and structural entropy identified in the cell cycle study) (**Fig. 4c**). We note that the fractometry strategy presented in this work could also be immediately applicable to the existing QPI modalities, especially because the imaging speed and throughput of the existing QPI modalities continue to increase to the scale comparable to those of flow cytometry ^51^.

As QPI can generally be adaptable with the typical fluorescence microscopes ^29,52^, we anticipate that single-cell fractometry could readily be incorporated into the current fluorescence-based morphological profiling strategies which are increasingly promising in many applications, from drug discovery to basic biology research ^53–55^. Using this multimodal imaging approach (especially in a high-throughput configuration ^56^), future studies could aim to systematically investigate how the biophysical fractal behaviours (i.e. mass fractal studied in this work) of different subcellular organelles (e.g. nucleus ^12^, mitochondria ^18^, and cytoskeleton ^19^) are influenced by chemical and even genetic perturbations. As the state-of-the-art single-cell computational tools become increasingly versatile in analysing not only traditional omics data (e.g. genomics, epigenetic, transcriptomic and proteomic etc.) but also cell morphological data ^57^, we anticipate that the inclusion of fractometry in morphological profiling could facilitate the discovery of the connections between multi-omics and cell fractal at single-cell resolution morphology, (especially how the molecular signatures dictates the disease-related fractal behaviors) – thus offering a new dimension for deciphering complex cellular heterogeneity.

## Methods

### Cell culture

Seven cancer cell lines were authenticated via the Human STR profiling cell authentication service, including three adenocarcinoma cell lines (ADC, H358 (EGFR WT), HCC827 (EGFR exon 19 del) and H1975 (L858R and T790M)), two squamous cell carcinoma cell lines (SCC, H520 and H2170) and two small cell lung cancer cell lines (SCLC, H526 and H69). The breast cancer cell line used for the cell cycle experiments was MDA-MB231. Leukemia cell lines used in this work were THP-1 (TIB-202TM) and Kasumi-1/ACC220 (CRL-2724TM). All cell lines used in this study were purchased from American Type Culture Collection (ATCC). Leukemia and lung cancer cell lines were cultured in the tissue culture flasks (surface area of 75cm^2^) (TPP) and MB231 in100mm culture dish (Labserv), which were both placed in a CO_2_ incubator with 5% CO_2_ under 37 °C. The full culture medium was ATCC modified RPMI-1640 (Gibco) supplemented with 10% fetal bovine serum (FBS) (Gibco) and 1% antibiotic–antimycotic (Gibco). Depending on cell confluency observed by a standard light microscope, passage or medium replacement was performed 2–3 times each week.

### Fluorescence labeling for cell-cycle tracking

Click-iT Plus EdU Flow Cytometry Assay Kits Alexa Fluor 488 and FxCycle PI/RNase Staining Solution were obtained from Invitrogen to define the ground-truth of the cell cycle stages (G1, S and G2 phases). The MDA-MB231 cell culture was firstly renewed for 8 mL medium mixed with 8 μL of 10 mM EdU staining solution. After 2-hour incubation, the cells were harvested by 0.25% Trypsin (Thermo Scientific) and washed by PBS with 1% BSA. The cells were then brought to protection from light for the following steps. The centrifuged cells were fixed by 100 μL Click-iT fixative (4% paraformaldehyde in PBS) and then permeabilized by the permeabilization and wash reagent (sodium azide) with 15-minute incubation. Next, 500 μL Click-iT Plus reaction cocktail was added into 100 μL cell suspension for 30 minutes under room temperature. After washing with 3 mL Click-iT permeabilization and wash reagent, the cell pellet was then mixed with 500 μL Click-iT permeabilization and wash reagent and 500 μL FxCycle PI/RNase staining solution for 30 minutes at room temperature. Lastly, after washing away the staining solution, PBS was added to make up a total volume of 7 mL cell suspension for the subsequent imaging experiments.

### Microfluidic channel fabrication

The channel was designed for optimization of inertial focusing to generate an in-focus single-cell stream under fast microfluidic flow (>2 m/s) and then fabricated by polydimethylsiloxane (PDMS) by soft lithography technique. Firstly, a layer of photoresist (SU-82025, MicroChem, US) was coated on a silicon wafer by a spin coater (spinNXG-P1, Apex Instruments Co., India), which was soft-baked for two times (65 °C for 3 minutes and 95 °C for 6 minutes). After cooling down under room temperature, a computer-aided design (CAD) pattern was transferred onto the photoresist by a maskless soft lithography machine (SF-100 XCEL, Intelligent Micro Patterning, LLC, US) through a 4-second exposure and a two-step post-baking (65 °C for 1 minute and 95 °C for 6 minutes). After photoresist development with SU-8 developer (MicroChem, US) for 5 minutes, the wafer was rinsed and dried for subsequent PDMS mixture pouring, which was mixed with PDMS precursor (SYLGARD^®^ 184 Silicone Elastomer kit, Dow Corning, US) and curing agent at a ratio of 10:1. The height control of the imaging section in the microfluidic chip was performed by placing a custom-designed acrylic block on the wafer, yielding a channel dimension of 30 μm in height and 60 μm in width. After the channel curing in an oven at 65 °C for 2 hours before demolding, a biopsy punch (Miltex 33-31 AA, Integra LifeSciences, US) was used to punch two holes for later tube insertion as the inlet and outlet of the channel. Afterwards, the bonding between the channel and glass slide was activated by oxygen plasma (PDC-002, Harrick Plasma, US) and oven baking under 65 °C for 30 minutes. Lastly, plastic tubings were inserted to the chip as channel inlet/outlet (BB31695-PE/2, Scientific Commodities, Inc., US).

### Multi-ATOM imaging

Multi-ATOM combines the time-stretch imaging technique^58,59^ and phase gradient multiplexing method to retrieve complex optical field information (including the bright-field and quantitative-phase contrasts) of the cells at high speed in an interferometry-free manner (**Fig. 1(a)**). Detailed working principle and experimental configuration were reported previously^8,25,26^. In brief, a wavelength-swept laser source was firstly generated by a home-built all-normal dispersion (ANDi) laser (centered wavelength: 1064 nm; bandwidth: ~10 nm; repetition rate: 11 MHz; pulse width = ~12 ps). The laser pulses were temporally stretched in a single-mode dispersive fiber (group-velocity dispersion (GVD): 1.78 ns/nm), and were then amplified by an ytterbium-doped fiber amplifier module (output power = 36 dBm with an on–off power gain = 36 dB). The pulsed beam was subsequently launched to and spatially dispersed by a diffraction grating (groove density = 1,200/mm) into a 1D line-scan beam which was projected orthogonally onto the cells flowing in the microfluidic channel. This line-scan beam was transformed back to a single collimated beam after passing through a double-pass configuration formed by a pair of objective lenses (N.A. = 0.75/0.8). Afterwards, the beam conveying phase-gradient information of the cell was split into 4 replicas by a one-to-four de-multiplexer, where each beam profile (*I_x_*^+^, *I_x_*^−^, *I_y_*^+^, *I_y_*^−^) was half-blocked by a knife edge from 4 different orientations (left, right, top and bottom) respectively. Recombining the 4 beams by a four-to-one fiber-based time-multiplexer, we were able to detect the line-scan phase-gradient information in 4 directions in time sequence at high speed by a single-pixel photodetector (electrical bandwidth = 12 GHz (Newport, US)). The digitized data stream was processed by a real-time field programmable gate array (FPGA) based signal processing system (electrical bandwidth = 2 GHz, sampling rate = 4 GSa/s) for primary cell detection and image segmentation with a processing throughput of >10,000 cells/s in real-time. These segmented phase-gradient images of cells were sent to four data storage nodes (memory capacity > 800 GB) through four 10G Ethernet links, which were reconstructed to 2D complex field information following a complex Fourier integration algorithm, detailed elsewhere ^25^ (**Fig. 1(b-c)**).

### Fluorescence detection

2-channel (i.e., 2-color) fluorescence detection was also synchronized with multi-ATOM, i.e., the bright-field, quantitative-phase contrasts and fluorescence signal from the same cell can be detected simultaneously. It was employed to generate the ground-truth of the cell-cycle stages for single-cell fractometry based on multi-ATOM images. When flowing through the imaging section in the microfluidic channel, the cells were excited by two continuous wave (CW) lasers (wavelength: 488 nm and 532 nm) simultaneously to generate epi-fluorescence signals, which were detected by two photomultiplier tubes (PMT) in the end. Frequency modulation was done (11.8 MHz and 35.4 MHz respectively) by a multichannel direction digital synthesizer to multiplex the PMT-received signals. After digital demodulation and low-pass filtering, the two fluorescence signals of each segmented cell were synchronized with multi-ATOM signal by the same FPGA configuration.

### Single-cell fractal analysis

The complex field at the image plane of the flowing cell measured by multi-ATOM is denoted as

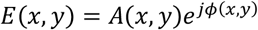

where *A*(*x*,*y*) is amplitude profile and *ϕ*(*x*,*y*)is the quantitative phase profile. In multi-ATOM, the phase gradient along the x-direction 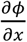 is first extracted based on the two raw knife-edged images (cut from left and right orientations): *I_x_^+^*(*x*,*y*) and *I_x_^−^*(*x*,*y*), through the relationship:

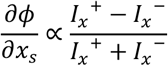

the same expression of the phase gradient along the y-direction 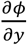 can be applied to the knife edges in the y-direction, *I_y_*^+^(*x, y*) and *I_y_*^−^(*x, y*). Hence, the quantitative phase *ϕ*(*x, y*) was then obtained by applying complex Fourier integration on the phase gradient images captured in multi-ATOM 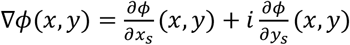:

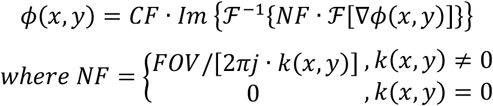

where *Im* is the imaginary part of a complex number; and 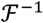 is inverse Fourier transform operator; *NF* is a normalization factor for quantifying the phase and avoiding singularity in the integration operation; *k*(*x, y*) is the 2D wavenumber; *FOV* is the 2D field-of-view; *CF* is the calibration factor for correcting the systematic phase deviation arise from non-ideal system setting^8^. On the other hand, the amplitude image of the cell (*A*(*x, y*)) is the sum of two images obtained from opposite knife edges normalized by the background (i.e., *B*, regions without samples).

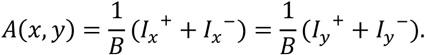

Subsequently, the complex field at the image plane is then numerically propagated to the far field using the Fourier transform operation – effectively yielding the (far-field) scattered light-field pattern 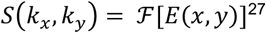, from which the fractal properties of cell can be measured. We then convert the scattered light pattern into an angular light scattering (ALS) profile *S*(*q*) in which scattered light intensity is averaged over rings of constant wave vector *q* = 4*π*/*λ*sin (θ/2), where *θ* is the *polar* scattering angle ^27^.

To quantify the fractal characteristics from the ALS, we also define a density-density correlation function in the real space, which is related to the ALS intensity via the Fourier transform relationship as the refractive index variation arises from density fluctuation *ρ*(*r*)^38^:

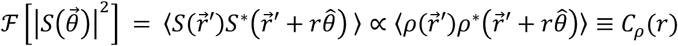

where *ρ*(*r*) means the density at point *r*, and thus *C_ρ_*(*r*) is the density correlation of an arbitary pair of occupied particles with a correlation distance of *r* ^60^. As It is known that the mass distribution of a fractal object can be expressed as ^9^, *m*(*r*) α *r^FD^*, where *m*(*r*) is the mass within a sphere of radius *r*. It can be also linked to *C_ρ_*(*r*) as ^60^:

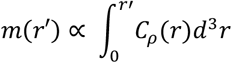

which indicates *C_ρ_*(*r*) ∝ *r*^*FD*–3^. Hence, the Fourier transform of an ALS profile will obey an inverse power law relationship, i.e., 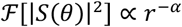, where *α* is the exponent, and FD can be expressed as:

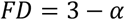

Therefore, by fitting the slope *α* of the log-scaled plot of 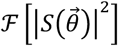 versus the correlation distance *r* we could quantify the FD, i.e., 3 – *α* (**Fig. 1(e)**).

## Supporting information

Supplementary for Morphological profiling by high-throughput single-cell biophysical fractometry

## Acknowledgements

This work is supported by the Research Grants Council of the Hong Kong Special Administrative Region of China (grant nos. 17208918, 17209017, 17259316; RFS2021-7S06, C7047-16G), and Innovation and Technology Support Programme (ITS/204/18).

## Author Contributions

K. K. T. conceived the project. Z. Z., K. C. M. L., D. M. D. S., Q. T. K. L. designed and performed experiments for fractal characterizations, with conceptual contributions from K. K. T. Z. Z. performed data analysis with interpretation from K. C. M. L., D. M. D. S., Q. T. K. L., and K. K. T. Z. Z. and K. K. T. wrote the manuscript, with assistance from K. C. M. L., D. M. D. S., Q. T. K. L., and K. K. Y. W. All authors edited the manuscript.

## Competing interests

The authors declare no competing interests.

## Notes

https://doi.org/10.5281/zenodo.6551588

### Competing Interest Statement

The authors have declared no competing interest.

