## Supplementary for Morphological profiling by high-throughput single-cell biophysical fractometry for "Morphological profiling by high-throughput single-cell biophysical fractometry"

### Table of content

Supplementary Table S1: Full list of Fourier-domain features.

Supplementary Fig. S1: Label-free single-cell image capture of 7 lung cancer cell lines.

Supplementary Fig. S2: Statistics of typical spatial-domain features of MB231 cell line.

Supplementary Fig. S3: 2-color fluorescence detection result by standard flow cytometry for cell cycle determination.

Supplementary Fig. S4: Feature performance on identifying the three cell cycle phases of MB231 cell line in one-versus-all mode.

Supplementary Fig. S5: Correlation matrix of all phenotypes of MB231 cell line (including both Fourier and spatial domain).

Supplementary Fig. S6: Circular plot combining the mean phenotypic heatmap and correlations of MB231 cell line with the feature labels.

**Table S1: Full list of Fourier-domain features.** 17 dimensions in total are extracted from ALS and correlation function to characterize the single-cell heterogeneity by multi-ATOM.

| Feature name | Physical meaning | Notation | Equation |
|---|---|---|---|
| FD | Fractal dimension | $FD$ | $FD = 3 - \alpha, [\alpha, \beta] = \frac{\boldsymbol{X}^T\boldsymbol{Y}}{\boldsymbol{X}^T\boldsymbol{X}}$ where $\boldsymbol{X} = \begin{bmatrix} \log(r_1) & 1 \\ \vdots & \vdots \\ \log(r_n) & 1 \end{bmatrix}, \boldsymbol{Y} = \begin{bmatrix} \log[f(r_1)] \\ \vdots \\ \log[f(r_n)] \end{bmatrix},$ $f(r) = F[S(\theta)]^2$, $n$ is the length of correlation function |
| FW Width | The width of fractal window | $r_j - r_i$ | Find the interval $[r_i, r_j]$ where $\frac{\log[f(r_k)] - \log[f(r_{k-1})]}{\log(r_k) - \log(r_{k-1})} < \alpha \ (i < k \leq j)$ |
| FD with FW | Fitted fractal dimension within the fractal window | $FD'$ | $FD' = 3 - \alpha', [\alpha', \beta'] = \frac{\boldsymbol{X}'^T\boldsymbol{Y}'}{\boldsymbol{X}'^T\boldsymbol{X}'}$ where $\boldsymbol{X}' = \begin{bmatrix} \log(r_i) & 1 \\ \vdots & \vdots \\ \log(r_j) & 1 \end{bmatrix}, \boldsymbol{Y}' = \begin{bmatrix} \log[f(r_i)] \\ \vdots \\ \log[f(r_j)] \end{bmatrix}$ |
| FD MSE1 | Fitting error within the fractal window | | $\frac{(\boldsymbol{X}'\boldsymbol{A}' - \boldsymbol{Y}')^T(\boldsymbol{X}'\boldsymbol{A}' - \boldsymbol{Y}')}{n}$ where $\boldsymbol{A}' = [\alpha', \beta']$ |
| FD MSE2 | Fitting error of the overall function | | $\frac{(\boldsymbol{XA} - \boldsymbol{Y})^T(\boldsymbol{XA} - \boldsymbol{Y})}{n}$ where $\boldsymbol{A} = [\alpha, \beta]$ |
| ALS | Angular light scattering (1D function) | $S(\theta)$ | $S(\boldsymbol{q}) = \int E(\boldsymbol{r})e^{-i\boldsymbol{q}\cdot\boldsymbol{r}}d^2\boldsymbol{r} = \mathcal{F}[E(\boldsymbol{r})]$, where $\boldsymbol{r}$ stands for the spatial vector, and $\theta = 2sin^{-1}(\frac{q\lambda}{4\pi})$ |
| ALS difference | The scattering difference of adjacent sample points (1D function) | $\Delta S(\theta)$ | $S(\theta) - S(\theta - \Delta\theta)$, where $\Delta\theta$ means the angular resolution of ALS measurement |
| ALS slope | The point-by-point slope of scattering curve (1D function) | $d(\theta)$ | $\Delta S(\theta)/\Delta\theta$ |
| ALS Drop – 5° | The intensity difference of scattering function from 0° to 5° | | $S(0°) - S(5°)$ |

| | | | |
|---|---|---|---|
| ALS Drop – 10° | The intensity difference of scattering function from 0° to 10° | | $S(0°) - S(10°)$ |
| First Minimal Height | The intensity difference of from 0° to first local minimal | | $S(\theta_0) - S(0°)$, where $\theta_0$ is the minimal value of $\theta$ to make $\frac{d}{d\theta}S(\theta_0) = 0$ and $S(\theta_0) \leq S(\theta_0 \pm \Delta\theta)$ |
| ALS Diff Peak | Maximum scattering difference | | $\max\,[\Delta S(\theta)]$ |
| ALS Slope Peak Drop | The scattering difference at the angle with the maximum slope | | $\max[d(\theta)] \cdot \Delta\theta$ |
| ALS Slope Peak | Maximum point-by-point slope | | $\max\,[d(\theta)]$ |
| ALS Mean | Mean of scattering function | $\bar{S}$ | $\mathrm{mean}[S(\theta)]$ |
| ALS Var | Variance of scattering function | $\sigma_S^2$ | $\sum_\theta [S(\theta) - \bar{S}]^2 / (N-1)$, where N is the length of ALS |
| ALS Skew | Skewness of scattering function | | $\frac{\sum_\theta [S(\theta) - \bar{S}]^3}{N \cdot \sigma_S^3}$ |
| ALS Kur | Kurtosis of scattering function | | $\frac{\sum_\theta [S(\theta) - \bar{S}]^4}{N \cdot \sigma_S^4}$ |
| ALS Range | Intensity range of scattering function | | $\max[S(\theta)] - \min\,[S(\theta)]$ |
| ALS Peak | Maximum value of scattering function | | $\max\,[S(\theta)]$ |

**Figure S1: Label-free single-cell image capture of 7 lung cancer cell lines.** Random-selected **(a)** phase gradient image in x-direction (DICx), **(b)** quantitative phase images (QPI) and **(c)** summarizing curve of the correlation function (ALS FT) of 7 lung cancer cell lines, respectively. Scalar bar is 5 µm. Shaded area indicates the statistical variance.

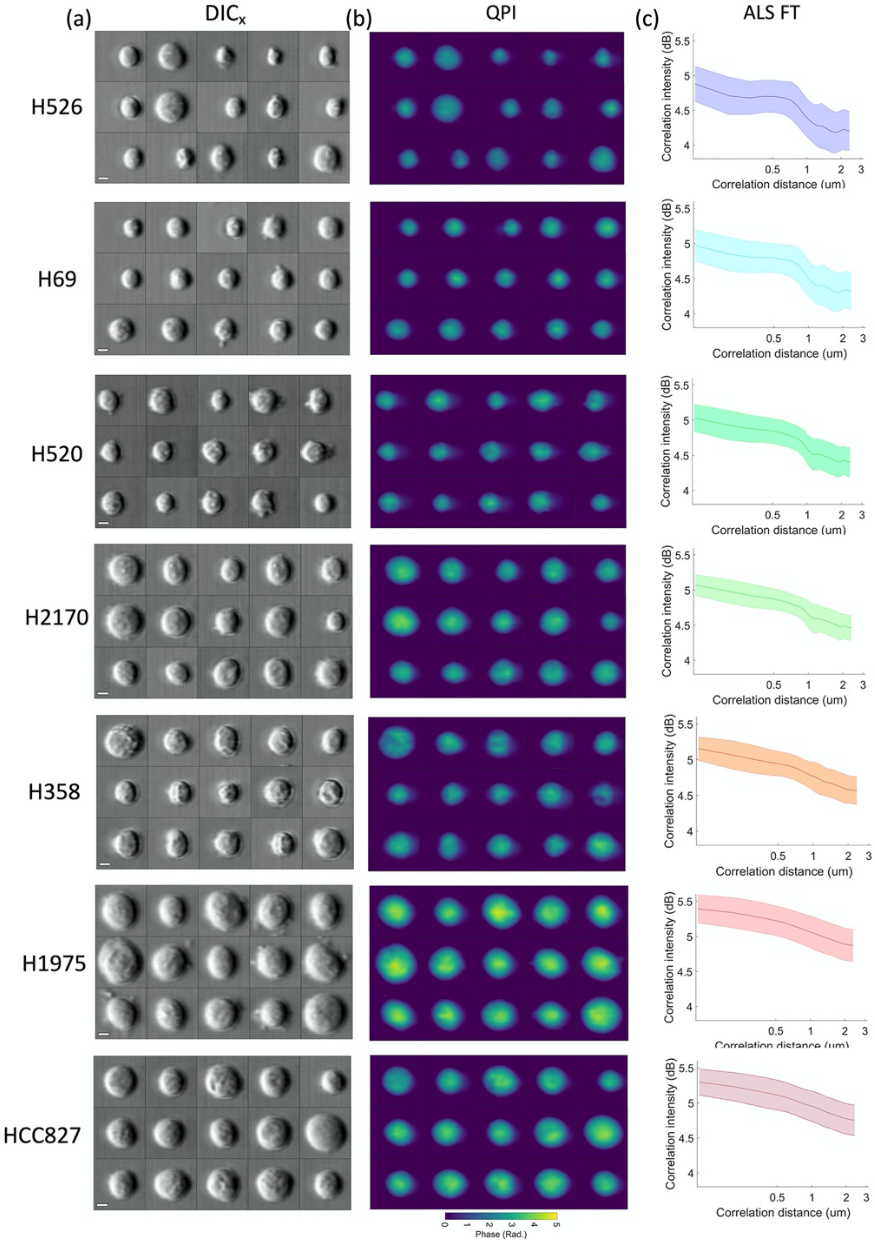

**Figure S2: Statistics of typical spatial-domain features of MB231 cell line.** Violin plot of **(a)** Cell size, **(b)** dry mass, and **(c)** dry mass density of cells in different cell cycle phase.

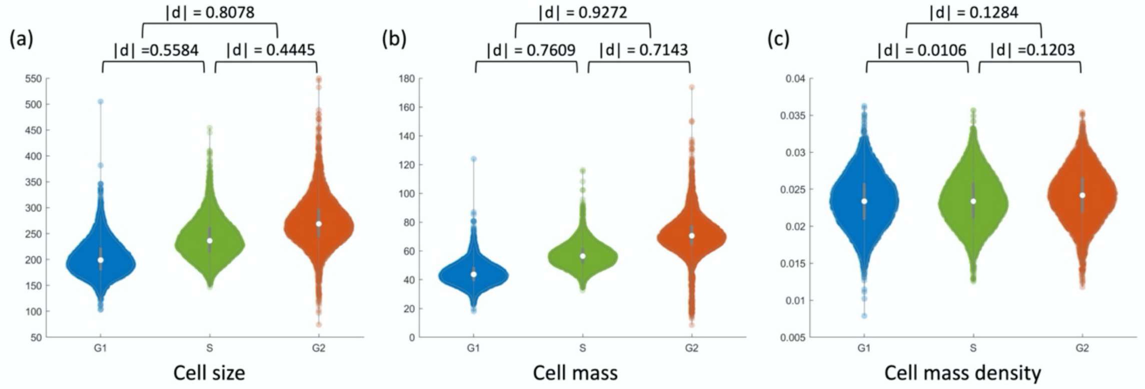


**Figure S3: 2-color fluorescence detection result by standard flow cytometry for cell cycle determination.** The cytometer used in this trail is *BD FACSAria™ III*. PI intensity is quantified in *PE-Texas Red* channel, whereas EdU intensity is detected in *FITC* channel. 50,910 cells are captured here in total, and 45,112 cells after gating of debris and aggregates possess a similar pattern showing cell cycle distribution with **Fig. 3b**.

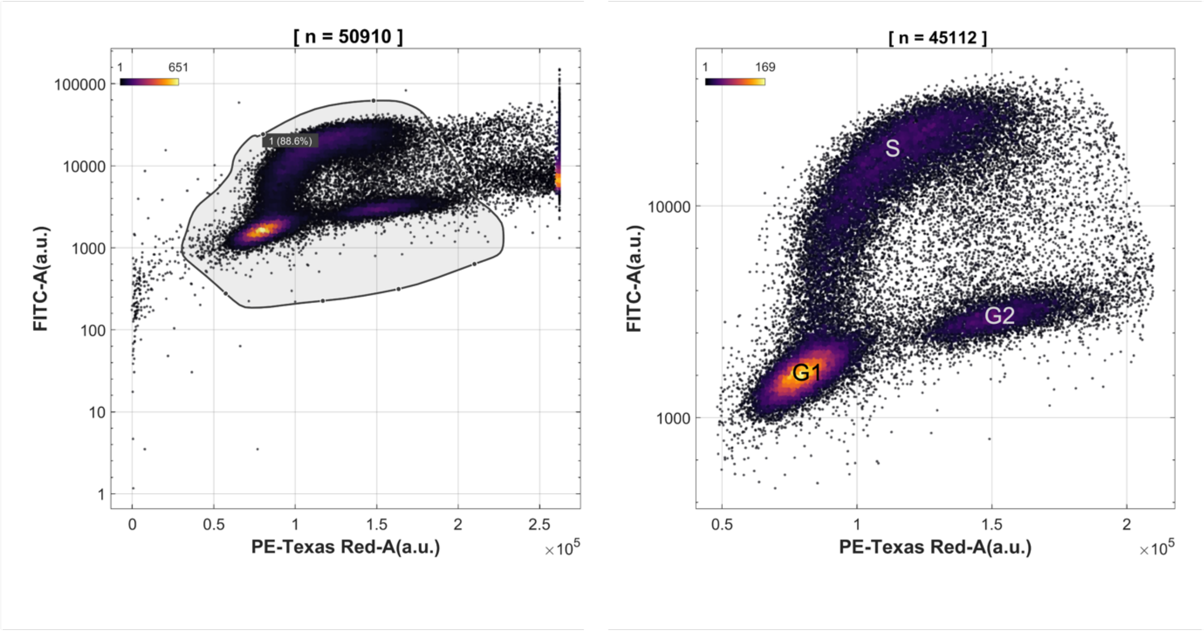

**Figure S4: Feature performance on identifying the three cell cycle phases of MB231 cell line in one-versus-all mode.** Features are ranked by the average area-under-curve of the receiver operating characteristics (AUROC) and only the top 15 features are shown.

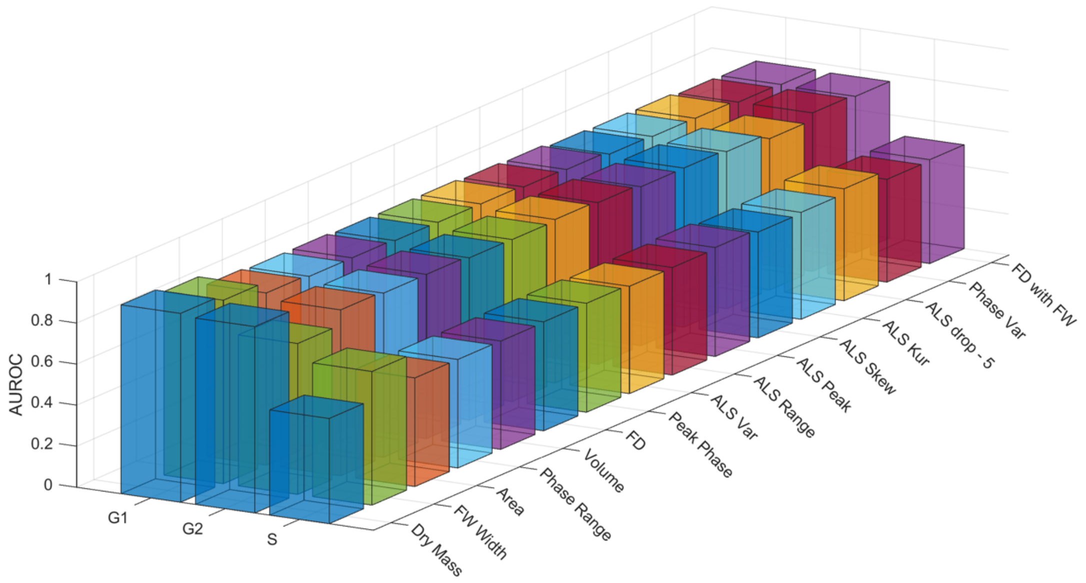

**Figure S5: Correlation matrix of all phenotypes of MB231 cell line (including both Fourier and spatial domain).** Blue and red color represents the positive and negative Spearman correlation coefficient, respectively, and the color level and size of the dot are both encoded with the magnitude of correlation coefficient. Features are arranged according to their feature type (i.e., bulk, global, local, fractal and ALS).

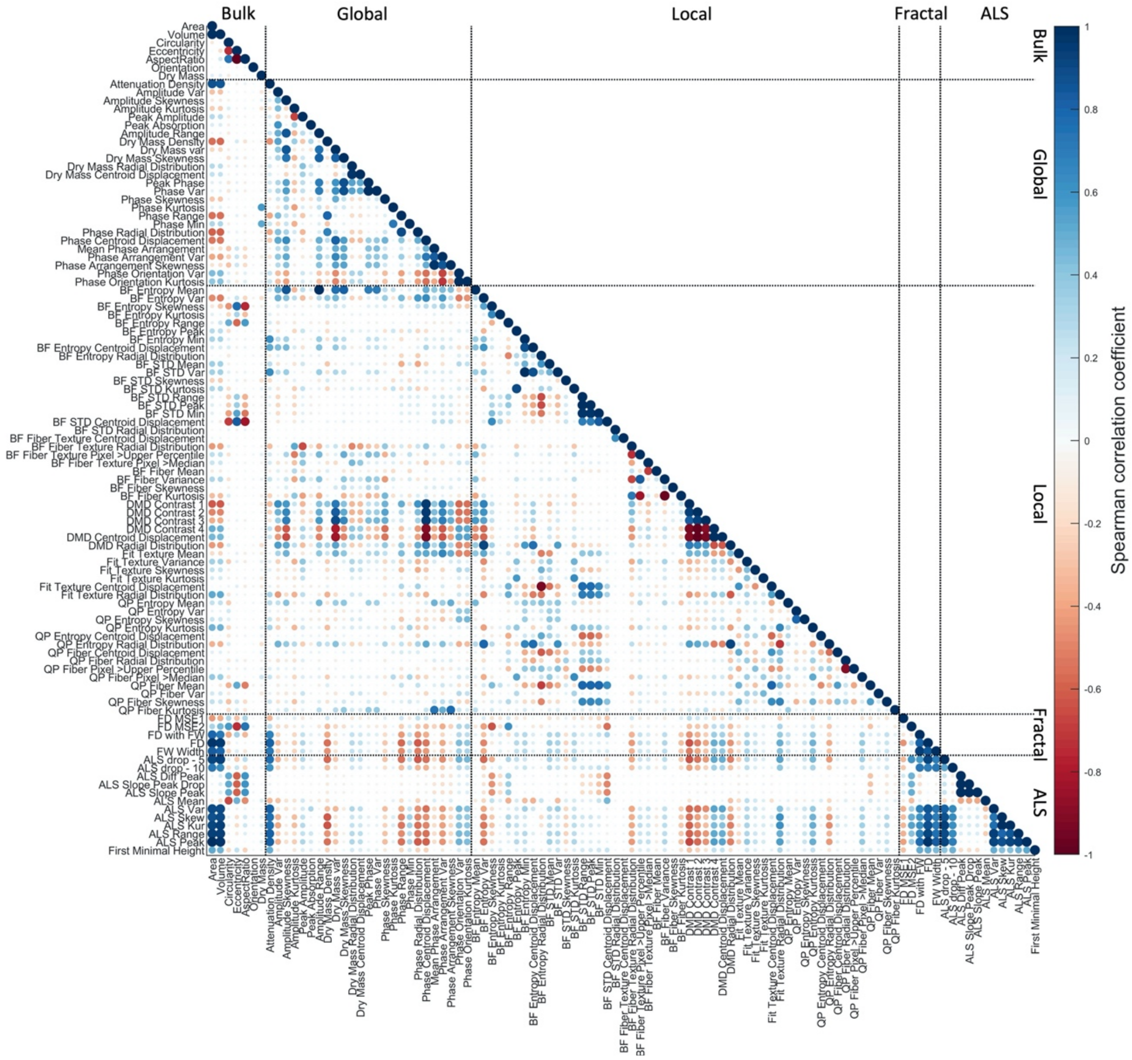

**Figure S6 (same as Fig. 4a): Circular plot combining the mean phenotypic heatmap and correlations of MB231 cell line with the feature labels.**

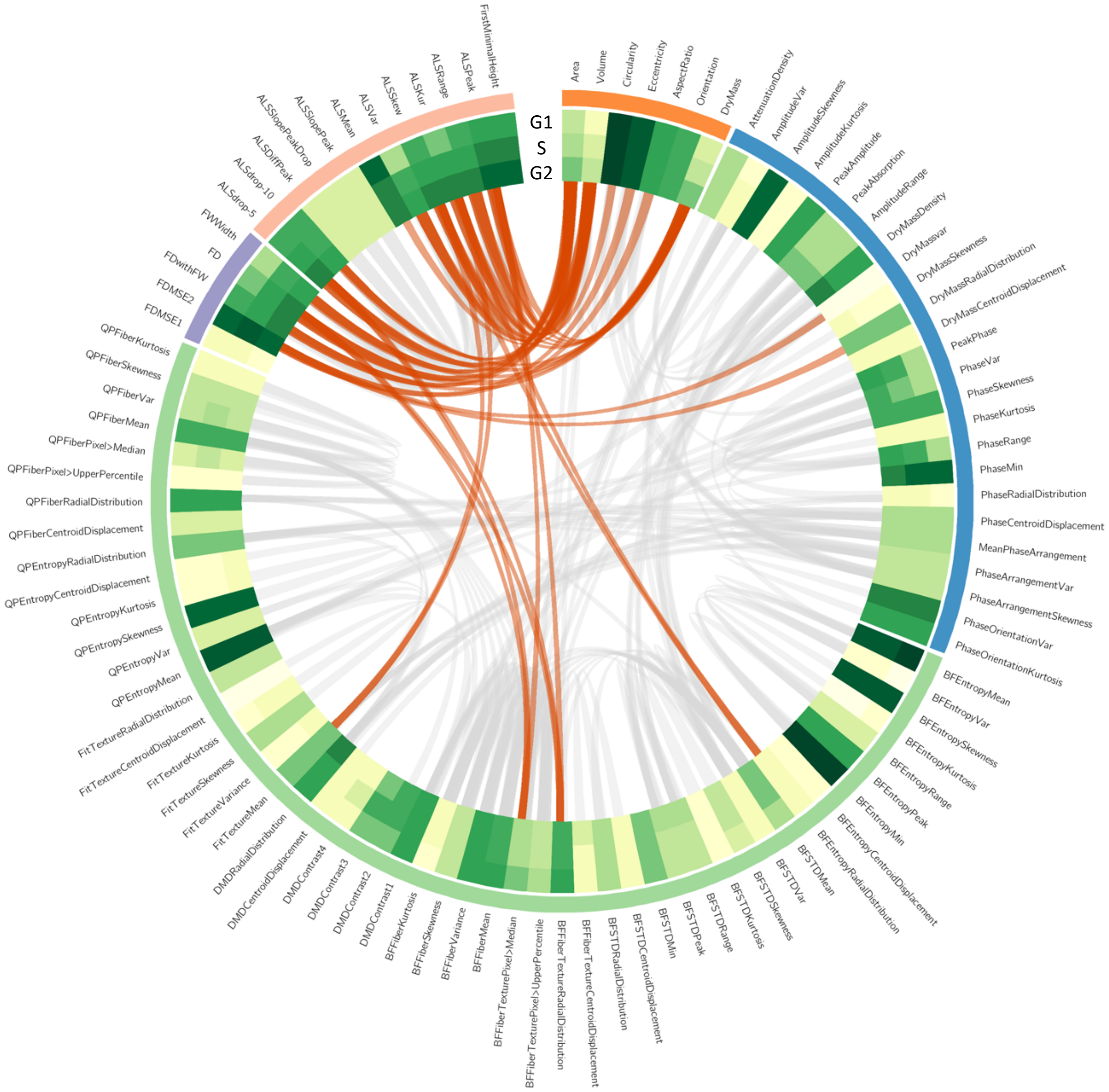
